# A mechatronic system for studying energy optimization during walking

**DOI:** 10.1101/496489

## Abstract

A general principle of human movement is that our nervous system is able to learn optimal coordination strategies. However, how our nervous system performs this optimization is not well understood. Here we design, build, and test a mechatronic system to probe the algorithms underlying optimization of energetic cost in walking. The system applies controlled fore-aft forces to a hip-belt worn by a user, decreasing their energetic cost by pulling forward or increasing it by pulling backward. The system controls the forces, and thus energetic cost, as a function of how the user is moving. In testing, we found that the system can quickly, accurately, and precisely apply target forces within a walking step. We next controlled the forces as a function of the user’s step frequency and found that we could predictably reshape their energetic cost landscape. Finally, we tested whether users adapted their walking in response to the new cost landscapes created by our system, and found that users shifted their step frequency towards the new energetic minima. Our system design appears to be effective for reshaping energetic cost landscapes in human walking to study how the nervous system optimizes movement.

## Introduction

Optimization is perhaps our most general principle of coordination. That is, people prefer to move in ways that minimize a cost function [1], [2]. A cost function is a weighted sum of one or more variables related to the movement, with the particular variables and their weights depending upon the task [1], [3]–[10]. For example, the cost function in reaching to a target is often modelled as a weighted sum of error and effort [6], [11]. One way to study optimization is to determine if the predicted cost minimum correlates with people’s preferences. Using reaching tasks as an example again, minimizing the sum of the variance about the target and squared muscle activations can explain people’s preferred arm trajectories [11]. While preferences suggest that the nervous system is concerned with optimizing coordination, preferences alone don’t provide insight into how it is accomplished. To study the nervous system’s optimization algorithms, it is useful to manipulate the values returned by the nervous system’s cost function and determine whether, and how, people respond. This is best accomplished in tasks where the cost function is well established, easy to manipulate, and directly measurable.

Walking is a well-suited task to study the nervous system’s optimization mechanisms. It has been well established that people typically prefer to walk in ways that minimize metabolic energetic cost [12]–[18]. For example, at every given speed, people choose to walk at the step frequency that minimizes their energy use [14], [15]. In addition to the many studies demonstrating preferences for energy minimal gaits in familiar conditions, recent research from our lab by Selinger *et al*. found direct evidence that the nervous system can continuously adapt these preferences to optimize energy during walking [19]. Selinger *et al*. used a knee exoskeleton to manipulate a user’s cost landscape by penalizing certain step frequencies, shifting the energy minimum away from the originally preferred step frequency. We use *cost landscape* to refer to the relationship between a gait parameter and its resulting metabolic energetic cost. Using such purposeful manipulation, Selinger *et al*. demonstrated that the nervous system could continuously optimize movements to converge on the new energy minimal gait within minutes [19]. Thus, energy optimization in walking is a useful model system because the nervous system’s cost function is unambiguous, we can measure the cost directly, we can directly manipulate the values returned by the cost function to study how the nervous system solves it, and the time course of optimization can be rapid enough that we can study it in a single experimental session.

This prior exoskeleton system allowed us to observe continuous optimization, but it was limited in its ability to probe the underlying mechanisms. The exoskeletons could only penalize users, and not reward them by decreasing their cost below their normal values. The range of penalties were limited to between 13% and 41% of the user’s original cost minimum, achieving a maximum gradient of 0.93% change in energetic cost for every 1% change in step frequency. In contrast, an ideal system would be able to precisely and rapidly prescribe cost landscapes of any shape. Such control over the shape of the cost landscape would allow both steeper and shallower cost gradients to study how the nervous system detects cost savings through its movement variability [20]–[22], the creation of complex cost landscapes to study the nervous system’s optimization algorithms [1], [23], and the lowering of energetic cost to study whether the nervous system differentially values energetic penalties and rewards [24], [25]. Applying new costs rapidly is also important because the nervous system’s association of a particular coordination pattern with its resulting energetic cost may depend upon the delay between the movement and its energetic consequence [26], [27]. A quick system can always be slowed down to study the effects of delayed costs, but an inherently slow system cannot be sped up.

Here we present a new mechatronic system designed to probe optimization mechanisms during walking. Based on data-driven simulations, the system uses controlled horizontal fore-aft forces applied near the center of mass of the user to change energetic cost landscapes. This system can generate a wide range of cost gradients because energetic costs during walking depend strongly on horizontal forces [28]. It can also provide both energetic penalties with backward forces and energetic rewards with forward forces. It uses a series elastic actuator to apply these controlled forces precisely and rapidly [29]. In the following sections, we first develop the high-level design using simulations that leverage literature data and then describe the construction of the system. Next, we evaluate system performance including a) how well it controls forces, b) how well the measured energetic costs match the predictions generated during design, and c) how users adapt their walking in response to new cost landscapes created by our system.

## System Design

We designed a system to manipulate the energetic cost of human walking. We achieve this by applying controlled horizontal fore-aft forces to the user as a function of one or more measured gait parameters. In this paper, we use a single parameter, step frequency, for ease of comparison with prior work [14], [19], [28]. For an individual walking at a constant speed, there is an energetic cost associated with each step frequency, with the minimal cost occurring at their preferred step frequency. We term this their *originally preferred step frequency*. Our system applies a backward or forward pulling force at each measured step frequency. Thus, the final energetic cost experienced by the user is the sum of the energetic cost associated with that step frequency and the energetic penalty (or reward) from the applied force. This new association between step frequency and energetic cost is the *new cost landscape*. In this manner, we can shift the energetic optimum higher or lower than the originally preferred step frequency. We then measure users’ preferred step frequency in the new cost landscape to determine their *new preferred step frequency*.

As a first test of our design, we simulated the system’s effect on a user’s energetic cost. We used literature data of the energetic cost associated with various walking step frequencies when no force is applied, where step frequency is measured as percent of the individual’s preferred step frequency [14]. We also used literature data for the energetic cost of walking at the preferred step frequency while a range of forces are applied, where the forces are measured as percent body weight of the individual [28]. We then combined these two data sets to obtain the energetic cost of walking at various step frequencies over a range of applied horizontal forces. Constraining the range of step frequencies to ±15% of preferred, and the range of horizontal forces to ±15% of body weight, we estimated that our system could vary a user’s energetic cost of walking from -45% to +230% relative to their original energetic minimum.

Based on these initial simulations, we built a mechatronic system to apply rapid, accurate, and precise forces as a function of the user’s measured step frequency (Fig. 1). In this system, users walk on a single-belt treadmill (Trackmaster TMX425C, Full Vision Inc., Kansas, USA) while wearing a hip-belt (Osprey Isoform4 CM) that places them in a closed loop with the actuator. The hip-belt is tailored with extended belt loops in the front and back to which we attach long inextensible nylon cables. The long lengths (~409 cm in front and ~197 cm in back) help ensure that the forces on the user remain nearly purely in the fore-aft direction despite the within-stride vertical and medio-lateral movements of the center of mass during walking. These cables pass through nylon pulleys with bearings (McMaster-Carr Nylon Pulley 3434T16) on either end and meet the actuator located behind the user (Fig. 1). We use light weight pulleys with low friction bearings to ensure low reflected inertia and minimal loss of forces during transmission. We use a series elastic actuator designed by Yobotics to produce the required force [29]. It consists of a 70 Watt brushless DC motor (BN23-23PM-03LH, Moog Inc.) that rotates a custom molded lead ball screw to maintain a set of four compression springs (McMaster-Carr Compression Spring 9434K147; spring stiffness = 33 lbs./in) at the required compression as determined by the commanded force. We measure the actual compression of the springs using a linear optical encoder (LIN-120-32, US Digital, Vancouver, WA, USA) and maintain their position with a proportional-integral controller implemented with a motor driver (Accelnet panel ACP-090-36, Copley Controls) that commands the motor using a 15 kHz center-weighted pulse-width-modulated signal. The required spring compression is commanded to the motor driver from a real-time controller (ds1103, dSPACE GmbH, Paderborn, Germany).

**Figure 1:**
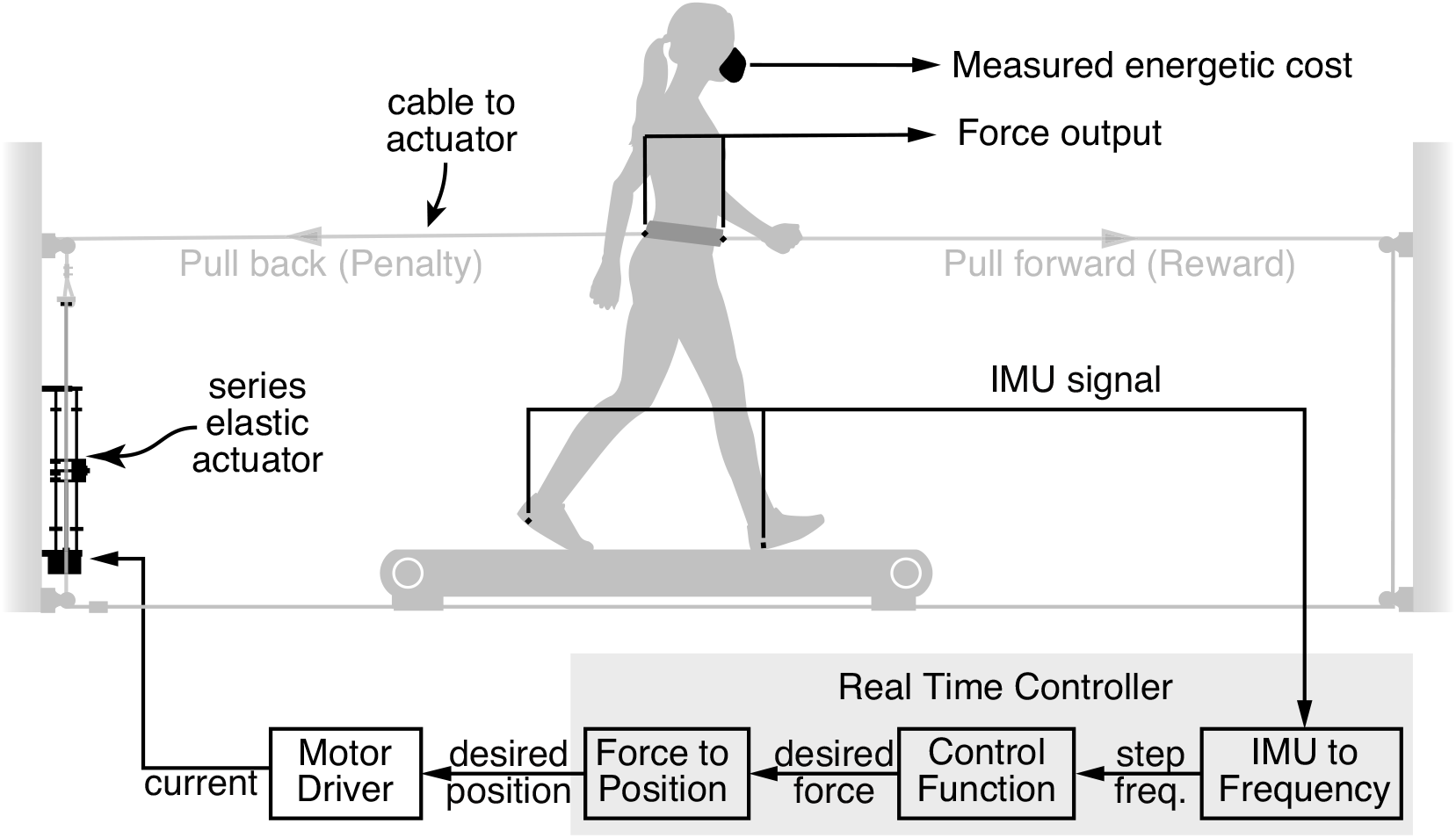
Mechatronic system. An actuator pulls backwards or forwards on a walking user via long tensioned cables attached to a hip belt. The cables are routed through pulleys, so that they attach on either side of the same linear actuator. Backward forces provide an energetic penalty, raising the cost of walking relative to normal. Moderate forward forces provide an energetic reward, lowering energetic cost. Force transducers between the cables and belt measure the forces applied to the user. Step frequency is measured from IMUs attached to the feet. Output forces from the motor are controlled using real-time custom designed control hardware and motor driver. Metabolic energetic cost is measured using indirect gas calorimetry.

This real-time controller measures step frequency, performs online calculations, and sends the resulting force commands to the motor driver. It is built in Simulink (Mathworks, NA, USA), and once compiled, runs at 1 kHz on the controller hardware. The controller receives analog signals from inertial measurement units (IMU) placed near the heel of each foot and estimates the step frequency by identifying foot ground contact events using characteristic zero crossings in angular velocity. To reduce error, it filters the analog signals using a 10 Hz, 2^nd^ order low-pass Butterworth filter and requires that consecutive zero-crossings arise from alternate feet. The implementation of this condition requires the controller to divide this value by two to obtain step frequency. It passes this step frequency through a *control function* that determines the force to be commanded depending on whether we want to reward or penalize the measured step frequency and by how much. We then use a pre-determined calibration function to estimate the spring compression required to produce the commanded force. The calibration function was obtained by measuring force output for a range of commanded spring compressions. Finally, the controller commands the estimated spring compression to the motor driver. We designed the controller to accept inputs from a MATLAB 2013b (Mathworks, NA, USA) script for parameters such as the user’s mass. We use dSPACE ControlDesk 5.2 (dSPACE GmbH, Paderborn, Germany) to monitor and record the data.

We also use the dSPACE board to monitor and record the forces and the signals required to calculate energetic cost measures.’We calculate the actual net force applied to the user as the sum of forces measured by the two load cells (LCM201, Omega Engineering) attached to the cables at the front and back of the user’s hip-belt. The forces are transmitted as analog signals to the dSPACE board where they are filtered using a 30 Hz, 2^nd^ order low-pass Butterworth filter. To calculate energetic cost, the dSPACE board records from oxygen, carbon dioxide, and flow sensors, mounted on the mask worn by the user (Vmax Encore Metabolic Cart, Viasys, Pennsylvania, USA).

## System Performance

We evaluated system performance at four levels. First, we considered open loop control of constant forces when there is no user present in the system. Second, we tested open loop control of constant forces while a user walked in the system. Third, we closed the force control loop using our controller and evaluated the control of forces when the commanded force changed with each walking step. Finally, we measured the effect of the system on the user’s energetic cost and gait adaptation. All walking trials were at a constant speed of 1.25 ms^-1^. All protocols with human users were approved by the Simon Fraser University’s Office of Research Ethics, and all users gave their written, informed consent before participation.

We first evaluated open-loop force control performance to determine whether our system could change between target forces within a walking step. We measured this responsiveness as *rise time*— time taken for the measured force to reach 90% of the way towards a new commanded value from an original commanded value. To accomplish this, we replaced the hip-belt with a wooden plank attached rigidly to the treadmill frame. We measured the force applied on the plank when we manually commanded step changes in force between 0N and either -49 N or +49 N. We repeated each step change 20 times, and each step lasted 4 seconds. We choose these force levels to match 10% of the body weight of our primary system tester, who happens to weigh 50 kg. By design, the magnitude of these step changes is conservatively large—when under closed-loop control, users would normally not make a large step-to-step adjustment in step frequency, and thus would not experience a step-to-step change in force as large as we tested here. We found an average rise time of 85 ms. Even assuming a high walking step frequency where each step takes only 400 ms, this easily allows for force changes within a single walking step [30](Fig. 2).

**Figure 2:**
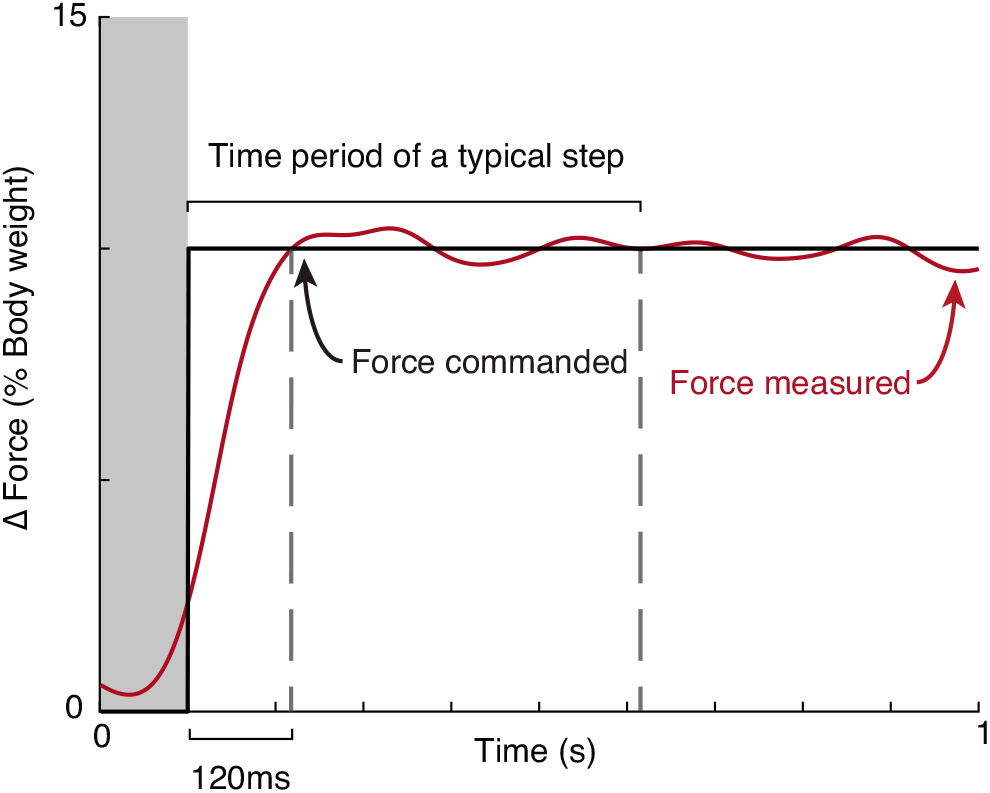
Open loop force control - Responsiveness. Time taken for the measured force (red) to reach a new commanded value from an original commanded value (black). Data shown here were averaged over 20 step changes. Each step was a change of 10% body weight—the original commanded value was always 0 N while the final value was ±10% body weight (body mass=50kg; acceleration due to gravity=9.81 ms^-2^).

We next evaluated the system’s accuracy and precision in applying the commanded force to a walking user. We quantified accuracy as the *steady state error* between the commanded force and the measured force and precision as the *steady state variability* in force about the steady state force. We measured the force applied to our system tester as they walked at their self-selected step frequency for two six-minute trials. During this time, we manually commanded step changes in force between 0N and ±10% body weight (body mass=50 kg, acceleration due to gravity=9.81 ms^-2^). Importantly, the force commands were in open-loop—they did not depend upon the user’s step frequency. Each condition lasted one minute, and each non-zero force was preceded by the zero force condition. We allowed 35 seconds for the self-selected step frequency to approach steady state [31], [32] and performed our analyses on the remaining 25 seconds. We first averaged the force over each step, obtaining a single force value per step, and then averaged this value over the 25 seconds. We determined steady state error as the difference between this averaged force and the commanded force for that condition. We found that our system can match a commanded force with an average steady state error of 0.13% body weight (Fig. 3). The average steady state variability for this user was 0.39% body weight when calculated between steps (root-mean-squared error). The force variability within a step was considerably higher at 2.64% body weight. We were not concerned about the magnitude of this within-step variability since our controller is designed to manipulate the energetic cost only at each new step since it applies force as a function of step frequency. Similarly, the exoskeleton of Selinger *et al*. [19] had considerable within step variability in the torque it applied to the knee. Because the applied horizontal forces affect energetic cost, any inaccuracies or imprecision in applied forces will result in inaccuracies and imprecision in the cost landscapes created by our system. Based on our modelling during system design and the identified force control performance described here, we predict steady state energetic cost errors of ~1.3% and steady state energetic cost variability of ~3.8%, relative to the average minimum energetic cost of regular walking [14]. One consequence of this step-to-step energetic cost variability is that it is necessary to average over several walking steps to accurately estimate the steady state energetic cost generated by the controlled forces. This is not a major practical concern for the experimenter as the significantly greater variability in breath-to-breath measures of energetic cost [33] normally requires averaging energetic cost over 2-3 minutes, or about 200-300 steps.

**Figure 3:**
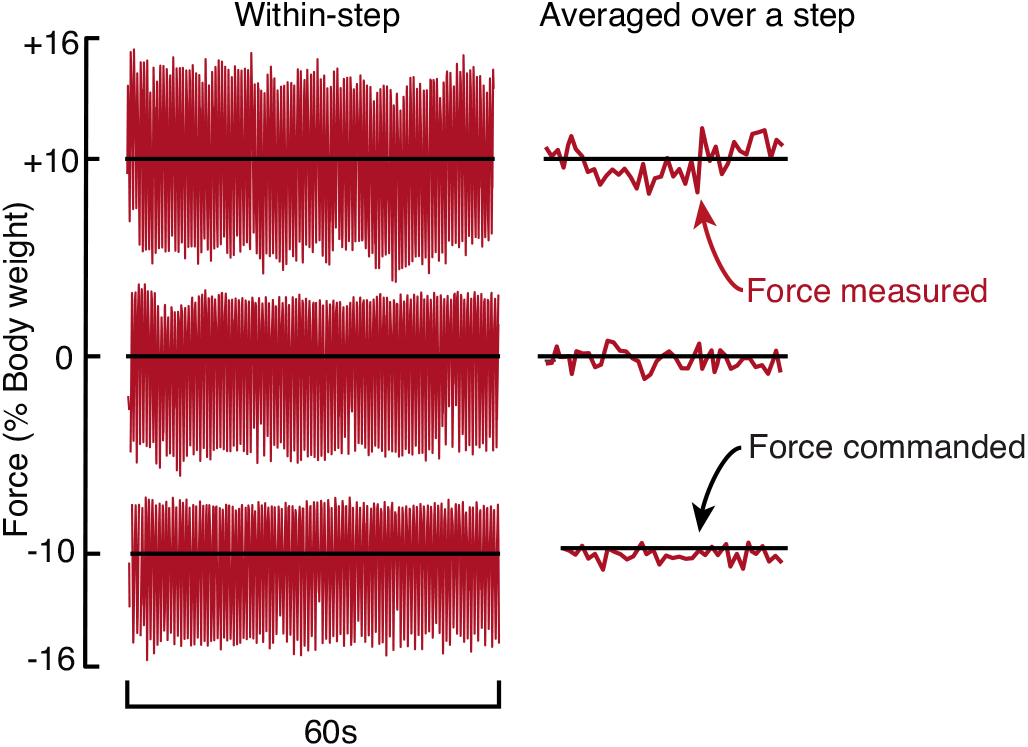
Open loop force control - Accuracy and Precision. The measured force (red) has a steady-state-error of 0.13% body weight when averaged over a step. The RMS error is 2.64% body weight within a step and 0.39% body weight when the force is averaged over a step. Data were collected from one user (body mass=50 kg; acceleration due to gravity=9.81 ms^-2^) walking at 1.25 ms^-1^ while attached to our system, with a constant force being commanded (black).

At the third level of evaluation, we applied the forces to a walking user as a function of their step frequency. We only evaluated precision at this level because there was no steady state force—the commanded force depended on the user’s execution of step frequency. We measured the net force applied to one user as they matched an audio metronome that played seven frequencies spanning ±15% of their originally preferred step frequency. Each step frequency condition lasted one minute, and the measured net force was first averaged over a step, and then over the last 25 seconds of that condition. We repeated this with two control functions where a control function defines the relationship between measured step frequency (*sf*) and commanded force (*F*):

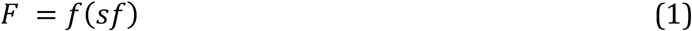

Both control functions were linear with zero offset, but one had a slope (*k*) of +1 thereby penalizing low step frequencies, while the other was -1 penalizing high step frequencies.

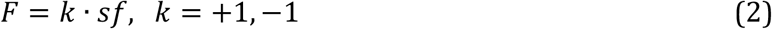

We averaged the steady state variability, calculated as root-mean-squared error, across all 14 trials and found it to be 0.59% body weight (user’s body mass=50 kg, acceleration due to gravity=9.81 ms^-2^) (Fig. 4). This results in a predicted steady state energetic cost variability of ~5.7%, which, as described in the previous paragraph, is not of practical concern for the experimenter when estimating steady state energetic cost.

**Figure 4:**
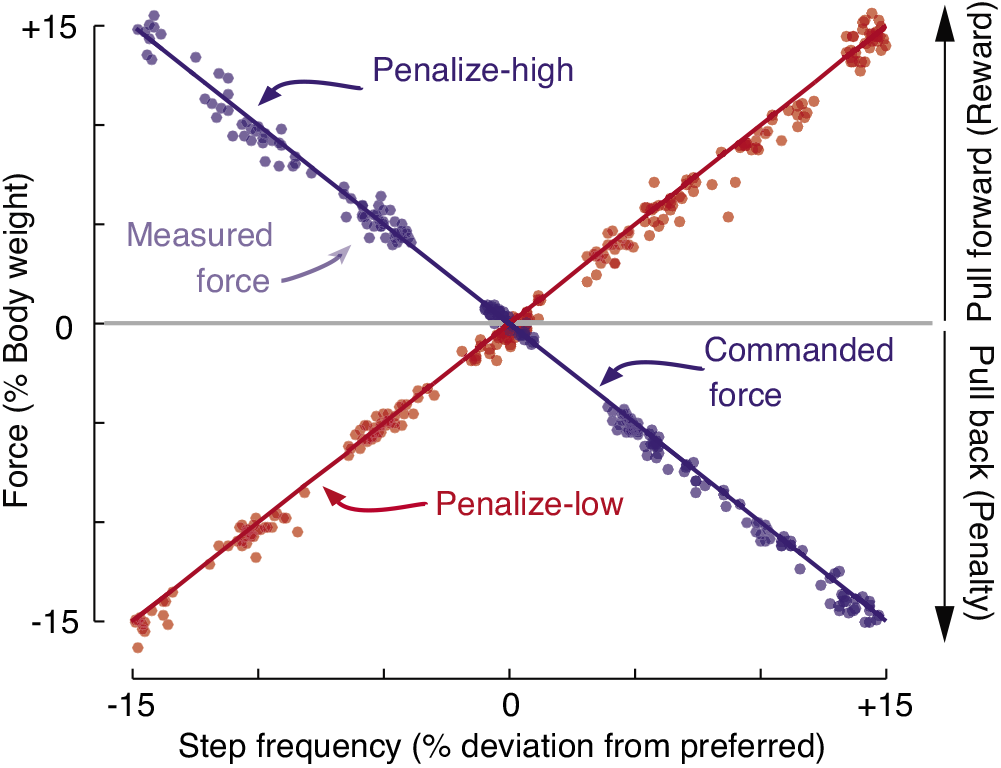
Closed loop force control. Measured force averaged over each step (dots) as a function of step frequency while walking at 1.25 ms^-1^ for a single trial by a single user. The user matched an audio metronome played at seven different step frequencies ranging from -15% to +15%. Red and blue lines illustrate that we used these forces to create two different cost landscapes—one that penalized low step frequencies (red), and the other that penalized high step frequencies (blue).

In the fourth level of evaluation, we created a new energetic cost landscape with our system and compared it to our predictions. Specifically, we measured how the metabolic energetic cost changed for one user as they walked in the system with a controller that commanded a force as a function of their step frequency. We tested the same two control functions described above. We designed them to create very steep cost landscapes that shifted the cost landscape minimum in different directions. First, we measured the user’s resting energetic cost during a 5-minute standing trial. Next, the user walked on the treadmill without the hip-belt, and unattached to the cables, for 12 minutes. We averaged the step frequency over the last three minutes to determine their originally preferred step frequency. Then, for each control function, the user walked at seven step frequencies spanning ±15% of their originally preferred step frequency. We enforced the step frequencies using a metronome and presented them in random order. To determine energetic cost, we measured the volume of oxygen consumed and volume of carbon dioxide produced using a respiratory gas analysis system (Vmax Encore Metabolic Cart, Viasys, Pennsylvania, USA). We divided these volumes by the measurement period to determine the rate of oxygen consumption 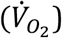 and carbon dioxide production 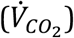. We applied the standard Brockway equation [34] to obtain the gross metabolic power:

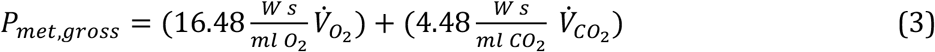

We define metabolic energetic cost as the energy used per unit time normalized for the person’s body mass (W/kg). This user’s body mass was 66 kg. Each trial lasted five minutes of which we allowed the first three minutes for the respiratory gases to reach steady state [35], and performed our analyses on the remaining two-minute measurement period. To compare with the user’s original cost of walking on a treadmill, we also measured their energetic cost at the same step frequencies while they walked on the treadmill without wearing the hip-belt or being unattached to the cables. We subtracted the user’s resting energetic cost for each condition and present here the net energetic cost. In accordance with our predictions, we could manipulate this users’ energetic cost of walking by as much as -49% to +230% of their original minimum (Fig. 5). This is more than a 5fold increase in the magnitude of applied penalty when compared to our exoskeletons, and unlike with our exoskeletons, our current system can provide an energetic reward by lowering cost.

**Figure 5:**
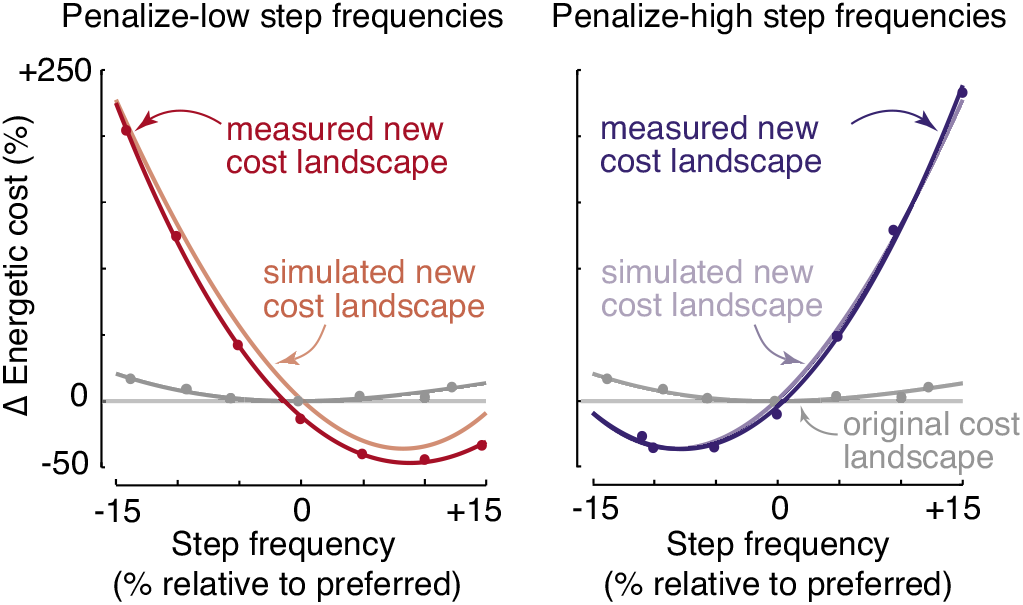
Simulated vs measured energetic costs. Metabolic cost measures (dots) for a single trial by a single user walking at 1.25 ms^-1^, when force changed as a function of step frequency as shown in Fig. 4. Grey curves illustrate the original cost landscape of the user while the red and blue curves illustrate cost landscapes that penalized low and high step frequencies respectively. Light colored curves illustrate our predicted results from simulations, and dark colored curves illustrate quadratic fits to measured data.

Next, we evaluated the ability of the nervous system to adapt gait towards the minima of new cost landscapes created by our system. To do so, we closely matched the cost landscapes and protocol used in our prior exoskeleton experiment, as that design was sufficient to generate continuous energy optimization during walking [19]. We used the following control function designed to penalize high step frequencies and shift the new cost landscape minimum to step frequencies lower than originally preferred:

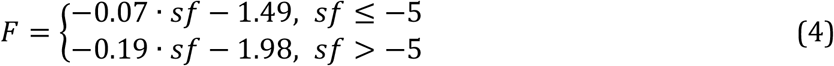

However, while prior studies parameterized the change in step frequency as percent deviation from the originally preferred step frequency, here we used step-to-step variability in step frequency, measured in standard deviations (SD). This allows us to understand the shift in preferred step frequency relative to the probability of it occurring by random chance. A 1SD shift was nevertheless comparable to a 1% shift as we found that the average standard deviation of our participants was 1.04%±0.12% (*N*=20).

Each participant completed three protocols on the same day. The purpose of the first protocol was to quantify each participant’s originally preferred step frequency, as well as the variability about the preferred step frequency (Fig. 6A). To accomplish this, each participant walked on the treadmill without the hip-belt for 12 minutes. To parameterize step frequency in future trials, we calculated the average and standard deviation of step frequency during the last three minutes.

**Figure 6:**
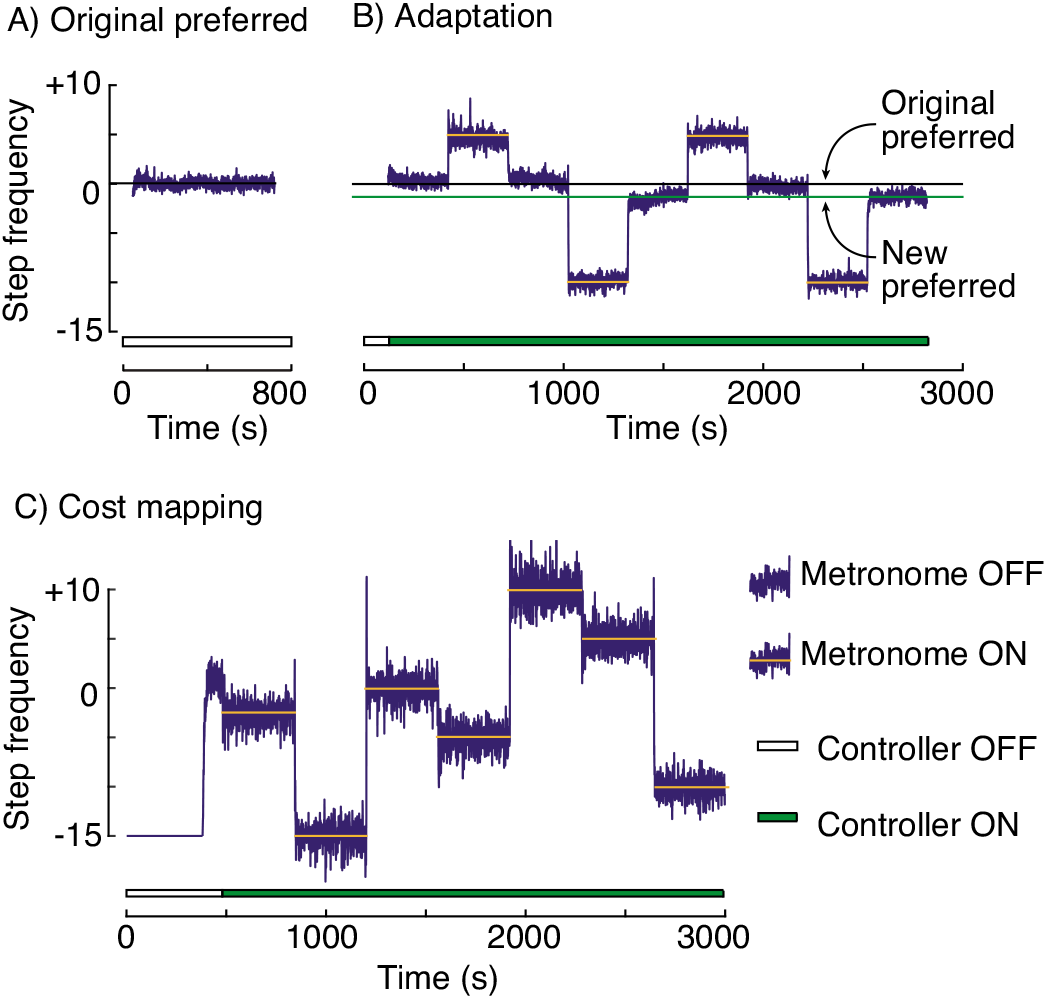
Experimental Protocol. A) and B) represent average measures from 8 participants while C) is from a single representative participant since the order of the metronome frequencies was different for each participant. **A)** Average originally preferred step frequency in the original landscape. **B)** Participants’ average step frequency time series during the protocol when we tested for gait adaptation. When the controller came on (green bar at the bottom) participants walked in a new cost landscape (Fig. 7) created by our system. Following the first five minutes of self-selected step frequency, they matched an audio metronome (orange bar) that either held them at a high step frequency (higher cost) or low step frequency (lower cost) relative to their originally preferred step frequency (black line) in the new cost landscape. C) Step frequency time series from a single representative participant during the third protocol when we measured their new cost landscape. Participants matched seven audio frequencies (orange bar) when walking in the new cost landscape (Fig. 7) while we measured their energetic cost.

The purpose of the second protocol was to determine the step frequency that participants preferred in the new cost landscape (Fig. 6B). To accomplish this, participants wore the hip-belt and walked continuously for 47 minutes while attached to our system. In the first 30 seconds, no force was applied, allowing the treadmill to reach the prescribed speed, and the participants to approach their originally preferred step frequency. Over the following 60 seconds, the force was slowly ramped up to match the force that participants experience at their originally preferred step frequency in the new cost landscape. This ensured that participants were not perturbed by a sudden force when we engaged the controller. The force was then held constant for 30 seconds, following which we engaged the controller, placing participants in the new cost landscape. Participants then alternated between periods of walking with a self-selected step frequency and walking to an audio metronome played at a prescribed frequency, with each period lasting five minutes. The metronome frequency alternated between +5SD and -10SD, to allow the participant to experience both higher and lower energetic costs relative to the cost of walking at their originally preferred step frequency in the new cost landscape. We determined participants’ new preferred step frequency in this new cost landscape as the average self-selected step frequency over the final three minutes of this protocol.

The purpose of the third protocol was to quantify the new cost landscape experienced by each participant (Fig. 6C). This was necessary since our hypothesis of observing adaptation towards the new energy optimum is conditional upon the new cost landscape having an energy optimum different from the original cost landscape. We first measured the energetic cost of standing still for six minutes for each participant. Next, participants walked in the system while under the same control function with the same force ramp up as described earlier. Participants matched seven step frequencies (-15SD, -10SD, -5SD, -2.5SD, 0SD, +5SD, and +10SD) produced by an audio metronome and played in random order. We measured each condition for six minutes. This was a minute longer than what we did with the previous control function (Eqn. 3) because we designed this cost landscape to be shallower. An extra minute of measurement provided us with more breath-by-breath samples of the energetic cost associated with each measured step frequency, and thus greater confidence in our cost estimates for each condition for each user. We subtracted resting energetic cost from each walking condition to obtain net energetic cost. Importantly, this was always the last trial that each participant performed, to ensure that it did not influence their gait adaptation.

Before analyzing our results, we tried to ensure that we had collected data from sufficient number of participants to effectively test for adaptation. We were primarily concerned with two issues. The first issue was that participants vary in the amount they shift their preferred step frequency in response to new energetic cost landscapes—the larger the variability, the greater the number of participants required to confidently detect a significant shift. To estimate the required number, we first needed to estimate the between-participant variability in self-selected step frequencies in new cost landscapes. To accomplish this, we conducted a pilot study with seven participants in a new cost landscape and found that the variability between participants in their self-selected step frequencies was 1.6SD. Using a power analysis for a one-tailed t-test, we determined that eight participants were required to detect a reduction of 2SD in their new preferred step frequency relative to their originally preferred step frequency (*β* > 0.90, *α* < 0.05).

The second issue was that individuals have different energetic responses to the same applied forces resulting in cost landscapes that vary between individuals. This variability, when combined with a control function designed to create small changes to the original cost landscape, results in some new cost landscapes that do not shift the new minimum cost to lower step frequencies. And we can only test the effectiveness of our system in causing adaptation to lower step frequencies when the new cost landscapes have a new energetic minimum at a lower step frequency. We operationalized this constraint by requiring that an individual’s new cost landscape meet three criteria—the cost at the new minimum, -2.5SD, and -10SD should all be more than 3SD lower than the cost measured at the originally preferred step frequency. We calculate this 3SD value from the variability in breath-to-breath energetic cost measured during the last three minutes of walking at 0SD. The first condition ensures that the new minimum provides a cost saving to the nervous system; the second ensures that there is a cost gradient at the originally preferred step frequency to direct the nervous system towards the new minimum; the third ensures that the nervous system experiences a cost saving during the metronome-enforced experience period. We performed a preliminary analysis of the cost landscapes of our first eight participants and estimated that we would need to collect from a total of 20 participants to have eight that met all three criteria. After collecting from 20 participants, we screened the cost landscapes and indeed found only eight that met the criteria. Importantly, we screened for these conditions prior to testing for gait adaptation in order to not bias our results towards a positive finding. Our subsequent analysis is of these eight participants.

Finally, we determined the step frequency that minimized energetic cost in the new cost landscape, and whether our participants adapted towards this new minimum. T o determine the step frequency that minimized cost, we used a linear mixed effects model to fit a quadratic relationship between step frequency and energetic cost to our measurements from all eight participants[36]. We chose this method, over fitting each participant’s costs individually, because breath by breath measures of energetic cost are quite variable, and this variability can dominate the differences in actual energetic cost between different step frequencies, causing noise to dominate when fitting individual cost landscapes. This method yields a value for the step frequency that minimizes energetic cost, but not the confidence in the location of this minimum. We used Monte Carlo simulations to determine the 95% confidence interval in this location. We resampled with replacement from the residuals of the linear mixed effects model and added it back to the average fit from the model. Each resampling yielded one simulated energetic cost for one step frequency for one simulated participant. We obtained 56 (8 participants by 7 step frequencies) such values to simulate one experiment and used a mixed effects model to fit these simulated values. We simulated 1000 such experiments and determined the location of the minimum in each case. We found that the average energetic minimum at - 5.7SD would have reduced the cost of walking by 6.1%, and the 95% CI of the location of the minimum spanned from -6.8SD to -4.8SD (Fig. 7). To compare, in the study by Selinger *et al*. with the exoskeletons, we observed energy optimization in new cost landscapes where the energetic minimum reduced the cost by 8.1% ± 7.0% (mean±SD) [19]. Using a one-tailed paired Student’s t-test, we found that in our new cost landscape, participants shifted their preferred step frequency away from their originally preferred step frequency and towards the energetic minimum by an average of -1.27SD (p=0.005) (Fig. 6B). This reduced their average energetic cost by 3.4%.

**Figure 7:**
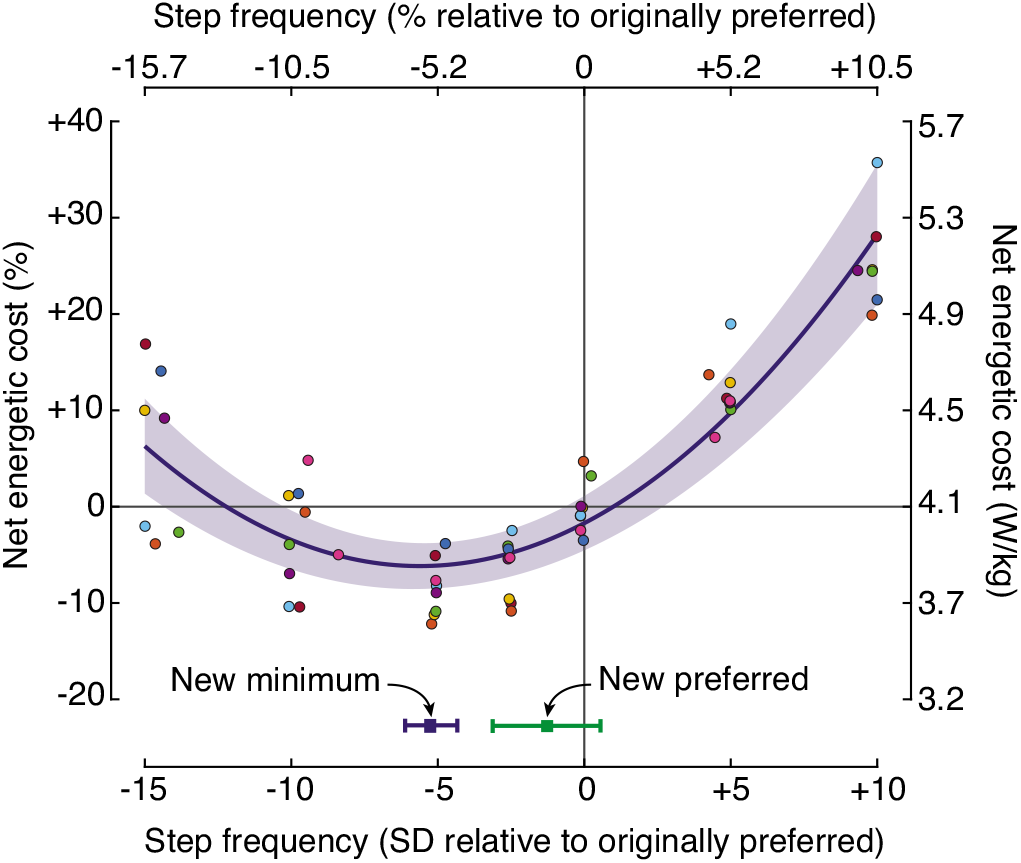
New cost landscape. The average new cost landscape where the energy optimal step frequency is shifted lower than the participants’ originally preferred step frequency (0SD). The green square shows the average and 95% CI of participants’ new preferred step frequency in the new cost landscape. This reduced their energetic cost by 3.4% relative to the cost their originally preferred step frequency. Each dot represents the steady-state energetic cost measured from a participant at the step frequency that they executed. Each color corresponds to one participant. The blue curve illustrates a 2^nd^ order linear mixed effects model of these data. The shaded region illustrates the 95%CI of the model, while the blue square at the bottom shows the minimum predicted by the fit, and the 95% CI of this minimum obtained using the Monte Carlo simulations. Left-hand y-axis values are normalized to the average energetic cost of 4.07 W/kg measured at 0SD.

## Discussion

Our system meets our performance requirements for studying energy optimization during walking. It can apply a large range of forces within the duration of a typical walking step. These forces are applied accurately and precisely even when the commanded force changes as a function of the user’s walking step frequency. These features together allow us to accurately manipulate the energetic costs associated with every walking step. The system can apply forces that both reward and penalize the energetic cost of walking, relative to the user’s original cost landscape. We can also accurately predict the average energetic cost landscape that the system creates for users for a given control function. The new cost landscapes created by our system caused users to adapt their walking gait towards the new energetic minimum, reducing their energetic cost by 3.4%. Interestingly, most users did not fully converge on the new optimum, which would have reduced their energetic cost by an additional 2.7% on average.

One candidate explanation for this incomplete energy optimization is that the optimization process is slow, particularly in our system. One factor that can slow optimization is measurement noise— the greater the variability in energetic cost, the greater the challenge for the nervous system to detect its gradient and assess the direction of the energetic minimum. This effect is especially pronounced near the minimum where the gradient is typically shallow, yet the noise is unchanged. The force variability in our system contributes to a noisy energetic cost gradient. For one user, we measured a between-step force variability of ~0.6% body weight resulting in a steady state energetic cost variability of ~6%. To confidently detect the gradient in the presence of this variability, the nervous system may need to average cost over longer periods of time, much like we do in our experimental methods, thus slowing down optimization. Indeed, learning tasks such as walking on a split-belt treadmill, crawling on hands and knees, and ergometer rowing can continue across many days of exposure [37]–[40].

Our system may be modified to reduce this force variability. We studied this in pilot experiments by first using closed-loop control of the forces applied to the torso, as measured with the force transducers, rather than the original design of maintaining a constant spring compression. We used proportional-integral-derivative control to drive the error between measured force and the commanded within-step force to zero. This appeared to successfully reduce the within and between step force variability, but we found that the controller gains were highly user-sensitive and required tuning specifically for each user. The tuning process required users to walk for ~10 minutes some of which time was spent feeling perturbed. This had several undesirable consequences including that users felt less safe in our system, that the nervous system had to discriminate between many different controllers in learning how to adapt, and that users were no longer naïve to the presence of a relationship between step frequency and the applied system forces. It is possible that these limitations may be overcome, but we decided to continue with the original approach at the possible expense of incomplete optimization. An alternative to closed-loop force control is to add compliance in series with the forward and backward pulling cables. We piloted this with three users and found them to adapt between 1SD and 3SD towards the new minimum, similar to the magnitude of adaptation we found in our original group of participants. While we suspect that the speed and completeness of optimization may increase with reduced force variability, we do not think it is necessary to eliminate it entirely. This is because not only was it present in the study by Selinger *et al*. where we did observe what appeared to be complete energy optimization, but such variability is also characteristic of everyday walking.

A second candidate explanation for incomplete optimization in our system is that it may be manipulating contributors to the nervous system’s cost function other than simply energetic cost. Our system applies physical forces to a user that change as a function of how they walk. It seems possible, if not likely, that our system changes the stability of walking along with energetic cost. We noted in the introduction that the nervous system’s cost function can be the weighted sum of one or more variables, with the particular variables and their weights depending upon the task. If stability normally contributes to the nervous system’s cost function during walking, or if the nervous system increases the contribution of stability to the cost function when walking in our system, the minimum of this cost function may not necessarily coincide with the minimum of our new energetic cost landscape—it may even be located at the step frequencies to which our participants adapted. Consistent with the possible contribution of stability to the cost function, pilot testing found that in steeper cost landscapes, the step to step changes in forces are perceptible and make walking uncomfortable. This means that while we can create cost landscapes of a wide range of gradients, and quickly apply these cost changes within a walking step, we may be required to trade-off between these two factors. We explored this in pilot experiments by designing a steep cost landscape but averaging step frequency over multiple steps before inputting into the control function that calculates the required force command. Since steep cost landscapes require large force changes between steps, the averaging reduces the effect these force changes have on stability by applying them over multiple steps. This may be an advantage to the nervous system for optimization as there are larger steady-state differences in energetic cost between steady-state step frequencies. Or, it may instead be a disadvantage as the nervous system may find it more difficult to associate variability in step frequency with the resulting changes in energetic cost. Thus, for the larger experiment we presented here, we chose to average step frequency only over a single stride and keep the cost landscape relatively shallow. We did design and pilot a cost landscape three times steeper than the one used here where the step frequencies were more heavily filtered through a three step mean filter. We tested two users and found that they were not noticeably uncomfortable with the force changes and shifted their step frequency by ~2SD towards the new minimum. This could be studied more systematically and is worth future research.

Were we to build our system again, there are several things we would change. First, we would consider using a different actuator. As described above, the series elastic actuator provided sufficient control of forces. But it was expensive, often required maintenance, and perhaps most importantly, it had limited travel. Due to this last limitation, we had to disqualify several pilot subjects who hit the limits while walking. We would replace the series-elastic actuator with a rotary motor, which would provide the system with unlimited travel. In our new design, the motor is housed off-board allowing one to use large, high-powered motors with high performance control of applied forces [41]. We suspect that this would allow us better control of the forces within and between steps. Second, we found that it was important to ensure a high degree of user comfort in the system. Towards this end, we would now prefer to use a wider and longer treadmill. This would help the user feel more comfortable when walking with potentially perturbing forces applied to them.

Our system, in its present form, is a robust tool for manipulating the energetic costs of walking in real-time. Here we focused on studying the performance of our device in healthy users that experienced shallow and simple cost landscapes which only penalized gait. Our system can also reduce the energetic cost of walking by as much as 50% allowing future studies to determine if the nervous system treats energetic rewards and penalties differently. The ability to create cost landscapes of various shapes will help study the nervous system’s optimization algorithms, including if they are speeded up by steeper cost gradients. Finally, reshaping cost landscapes based on rehabilitation goals may allow the nervous system’s internal drive to reduce energetic cost to aid gait rehabilitation in patients recovering from injuries and disorders [42].

## Acknowledgements

We thank Brian Umberger for providing the data we used to simulate the relationship between energetic cost and step frequency, Jessica Selinger for early input into the system and study design, and the SFU Locomotion Lab for their constructive feedback on manuscript drafts. This work was supported by a Discovery Grant from the Natural Sciences and Engineering Research Council of Canada and a US Army Research Office grant (W911NF-13-1-0268).

